# The missing role of gray matter in studying brain controllability

**DOI:** 10.1101/2020.04.07.030015

**Authors:** Hamidreza Jamalabadi, Agnieszka Zuberer, Vinod Jangir Kumar, Meng Li, Sarah Alizadeh, Ali Amani Moradi, Christian Gaser, Michael Esterman, Martin Walter

**Affiliations:** Department of Psychiatry and Psychotherapy, University of Tübingen, Tübingen, Germany; Boston University School of Medicine, Department of Psychiatry, Boston, USA; Boston Attention and Learning Laboratory, VA Boston Healthcare System, Boston, USA; Clinical Affective Neuroimaging Laboratory, Magdeburg, Germany; Leibniz Institute for Neurobiology, Magdeburg, Germany; Max Planck Institute for biological cybernetics, Tübingen, Germany; Department of Psychiatry and Psychotherapy, Jena University Hospital, Jena, Germany; School of Engineering, RMIT university, Melbourne, Victoria, Australia; Neuroimaging Research for Veterans Center (NeRVe), Veterans Administration, Boston Healthcare System, Boston, USA; National Center for PTSD, VA Boston Healthcare System, USA

**Keywords:** Network control theory, Gray matter, brain controllability

## Abstract

Brain controllability properties are normally derived from the white matter fiber tracts in which the neural substrate of the actual energy consumption, namely the gray matter, has been widely ignored. Here, we study the relationship between gray matter volume of regions across the whole cortex and their respective control property derived from the structural architecture of the white matter fiber tracts. The data suggests that the ability of white fiber tracts to exhibit control at specific nodes not only depends on the connection strength of the structural connectome but additionally strongly depends on gray matter volume at the host nodes. Our data indicates that connectivity strength and gray matter volume interact with respect to the brain’s control properties, such that gray matter exerts the great impact in regions with high structural connectivity. Disentangling effects of the regional gray matter volume and connectivity strength, we found that frontal and sensory areas play crucial roles in controllability. Together these results suggest that structural and regional properties of the white matter and gray matter provide complementary information in studying the control properties of the intrinsic structural and functional architectural of the brain.

## 2 Introduction

Network control theory, as recently applied to white matter fiber tracts in the human brain, provides a novel mechanistic framework to describe the ease of switching between different dynamical functional brain states, and the regions that best drive these dynamics (Bassett and Sporns, 2017; Medaglia, 2019; Medaglia, et al., 2017a). This approach has the potential to inform theories of dynamic cognitive processes, clinical neuroscience, neurodegeneration, and brain reserve. Specifically, there is evidence that these global brain state transitions are impaired in clinical populations (Braun, et al., 2019; Jeganathan, et al., 2018; Kenett, et al., 2018) and that such impairments can be traced back to specific driver nodes (Jeganathan, et al., 2018; Kenett, et al., 2018; Muldoon, et al., 2016; Zoeller, et al., 2019). However, this far, these control properties have been exclusively derived from white matter fiber tracts without the consideration of gray matter properties. Given the importance of GM properties for cognitive functioning and brain health, and the established interrelationships between white and gray matter, it has been suggested that regional gray matter integrity may be a critical contributor and proxy for network and node controllability (Medaglia, et al., 2017a; Medaglia, et al., 2017b).

Several lines of research suggest that GM may be essential to understanding brain controllability. First, GM is a proxy for the quantity of neurons and synaptic densities in a particular region (Lüders, et al., 2002), and metabolic energy expenditure is primarily realized through the gray matter cell bodies that scaffold white matter tracts (Zhu, et al., 2012). In neurodegenerative disorders, lesions of GM of a specific region only partially go along with corresponding lesions in WM in some neurodegenerative disorders (Agosta, et al., 2011; Bodini, et al., 2009; Douaud, et al., 2007; Raine, et al., 2000; Villain, et al., 2008), suggesting that GM reserve and WM provide independent/additional information with respect to controllability properties of the structural connectome. Taken together, these studies motivated the hypothesis that the controllability properties suggested by the WM should be partially related to or even predicted by GM integrity. Critically, it has been argued that including GM metrics in control theory will extend traditional volumetrics into network neuroscience (Medaglia, et al., 2017a). Nevertheless, to our knowledge the nature of the interdependency between controllability properties and GM properties has not been addressed empirically.

To tackle this issue, we used two independent data sets to investigate whether, and if so, how control properties extracted from the structural connectome relate to properties of the gray matter, i.e. GM volume which engenders other GM metrics e.g. surface and thickness (Kong, et al., 2015; Winkler, et al., 2010). Since previous studies have shown that brain controllability can be largely explained by the connectivity strength of the structural connectome, we also considered whether GM volume could explain additional variance in controllability not accounted for by white matter connectivity. Initially, we investigated how WM and GM factors affect brain controllability on a global brain level. In a further step, we identified the brain regions for which controllability was most sensitive to GM and/or WM properties. We discuss our findings with respect their potential relevance to cognitive and clinical neuroscience.

## 3 Methods and Materials

### 3.1 Data acquisition

The structural and diffusion datasets were taken from the Human Connectome Project (HCP, Principal Investigators: David Van Essen and Kamil Ugurbil; 1U54MH091657; Van Essen, et al., 2012). The data were selected from the 65 healthy subjects with the age range of 22 to 36 (28 M, mean age 29.2).

#### 3.1.1 MRI Data Specification

Structural images were acquired with the following specification: T1w MPRAGE, TR 2400 ms, TE 2.14 ms, TI 1000 ms, flip angle 8 degrees, Field of View (FOV) 224×224, 256 slices, voxel size 0.7 mm isotropic, Bandwidth 210 Hz/Px, IPAT 2, acquisition time 7:40 min.

DWI data were acquired by using a Spin-echo EPI sequence with TR 5520 ms, TE 89.5 ms, flip angle 78 degrees, voxel size, 1.25 mm isotropic, 111 slices, multiband factor, 3, echo spacing, 0.78 ms, b-values 1000, 2000, and 3000 s/mm2. For details, see (Glasser, et al., 2013; Van Essen, et al., 2012).

#### 3.1.2 AAL mask definitions

The 3-D anatomy atlas of the AAL was acquired from the neurofunctional imaging group (http://www.gin.cnrs.fr/en/tools/aal-aal2/) (Tzourio-Mazoyer, et al., 2002).

#### 3.1.3 AAL Native space transformation

The 12-parameter affine transformation (Jenkinson, et al., 2002; Jenkinson and Smith, 2001) was computed for each volunteer’s T1 and non-diffusion image and the MNI spaced standard brain. The resulted transformation matrix was applied to the left and right AAL brain regions to transform them into the native structural and diffusion space.

#### 3.1.4 Structural volume analysis

The tissue type segmentation employed SPM12 unified segmentation approach. The process resulted in segmented gray, white, and cerebro-spinal fluid (CSF) volumes. In the next step, we determined the volume of the brain, gray matter, and under each AAL atlas region for all subjects.

#### 3.1.5 Preprocessing and Diffusion-Fit

The obtained HCP diffusion data were reconstructed by using a SENSE1 algorithm (Sotiropoulos et al. 2013). The DWI data was corrected for motion and distortion (Andersson et al. 2003; Andersson and Sotiropoulos 2015, 2016). Furthermore, pre-processing included unringing, denoising, and tensor analysis implemented in MRtrix (Tournier, et al., 2012).

The data were reconstructed by using the multi-shell multi-tissue constrained spherical deconvolution (Jeurissen, et al., 2014). The resulted Orientation Distribution Function (ODF) was registered to the structural space. The initial tractogram was generated using mrtrix-tckgen, resulting in 100 million streamlines within each subject. In the next step, we applied spherical deconvolution informed filtering of tractograms (SIFT) to reduce the streamline count to 10 million. In the final step, the number of streamlines was determined between AAL brain regions to produce a connectome.

### 3.2 Network control framework

Controllability is one of the fundamental concepts in the mathematical control theory. The notion of controllability of a dynamical system was first introduced in (Kalman, 1963). State (output) controllability of a dynamical system is defined as the possibility of driving states (outputs) of the system from an arbitrary initial condition to any desired values in a finite time by applying appropriate control signals (Kailath, 1980). The most famous classic method to ensure state controllability of a dynamical system defined by the noise-free linear continuous-time and time-invariant network model says that the system

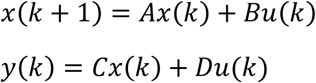

is full state controllable if and only if the Kalman’s controllability matrix [*B, AB*, …, *A*^*n*-1^*B*] has full rank (Kailath, 1980). ***x*** ∈ ℝ^*n*^ and ***u*** ∈ ℝ^*p*^ are state and input signals, respectively and *A, B, C* and *D* are matrices with appropriate dimensions where *A* and *B* are called state and input matrices, respectively. When applied in the context of brain controllability, ***x*** describes the state of brain regions over time, and A is an adjacency matrix. *A* determines the interaction between brain elements and its elements often represent the weight of the white tract connecting two areas (see section 3.3 for details). The input matrix **B** identifies the control nodes in the brain which may be confined to one node or a brain area or any combinations of nodes available in ***x***.

There are important relationships between controllability and stabilizability of both linear finite- and infinite-dimensional control systems. A linear dynamical system is stabilizable if its unstable modes are controllable (Chen, 1998). Controllability is also strongly related to the theory of minimal realization and canonical forms of dynamical system (De Schutter, 2000; Hazewinkel and Kalman, 1976; Rissanen, 1974). The minimum energy control problem for many classes of linear and time-delay systems is also related to the concept of controllability (Klamka, 2013). The positive semidefinite matrix

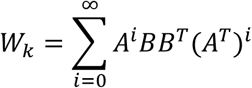

is controllability Gramian in which, eigenvectors associated with the smallest eigenvalue define directions in the state space which are less controllable and vice versa for eigenvector related to the largest eigenvalue (Kailath, 1980).

Although control theory is a mathematically highly developed branch of engineering, fundamental questions, raised in the recent decades, pertaining to the controllability of complex networks makes applicability of traditional approaches questionable (Liu, et al., 2011). Traditional controllability conditions should be updated considering a new player: Network topology. In addition, computationally efficient measures are required to study of real-world large size complex networks. These difficulties have been the root of recent research in controllability of complex networks. A promising approach has been to use the properties of controllability Gramian to assess the control properties of the brain when controlled from one single node. To perform the analysis, we chose average controllability (AC) which is the most commonly used controllability measure in the neuroimaging literature (Gu, et al., 2015; Medaglia, 2019). AC is defined by the trace of the controllability Gramian *Tr*(*W*_*k*_) and represents the average energy from each node when that node steers the brain into all possible output states. This value shows on average how much each node can move the brain in different directions which is a measure of ease of state transition capacity as reflected in the energy expenditure of the brain control system.

### 3.3 Statistical analysis: mixed effect model

To predict AC based on structural measures of the brain we built a linear mixed-effects (LME) regression (Baayen, et al., 2008) using a step-wise approach retaining an effect only if there was a significant difference between the log-likelihood ratio of the two models, based on an ANOVA (p < 0 05). Statistical analysis was performed using the lme4 package in R (Bates, et al., 2014).

## 4 Results

### 4.1 Effects of gray matter on average controllability

In a first step, we investigated if we could replicate previously reported findings that higher nodal degree (connectivity strength of a node) relates to higher AC (see Figure 1A). We also included the volume of the entire brain as a covariate, which has shown to be a confound of several properties of the GM (Lüders, et al., 2002). We built a linear mixed effects model to predict AC based on nodal degree with subjects as a random intercept. Our results replicate previous findings (Gu, et al., 2015) suggesting that structural connectivity strength across the whole brain increases global AC (β= .97, .88 – 1.06, p < .001) see Figure 1A).

**Figure 1:**
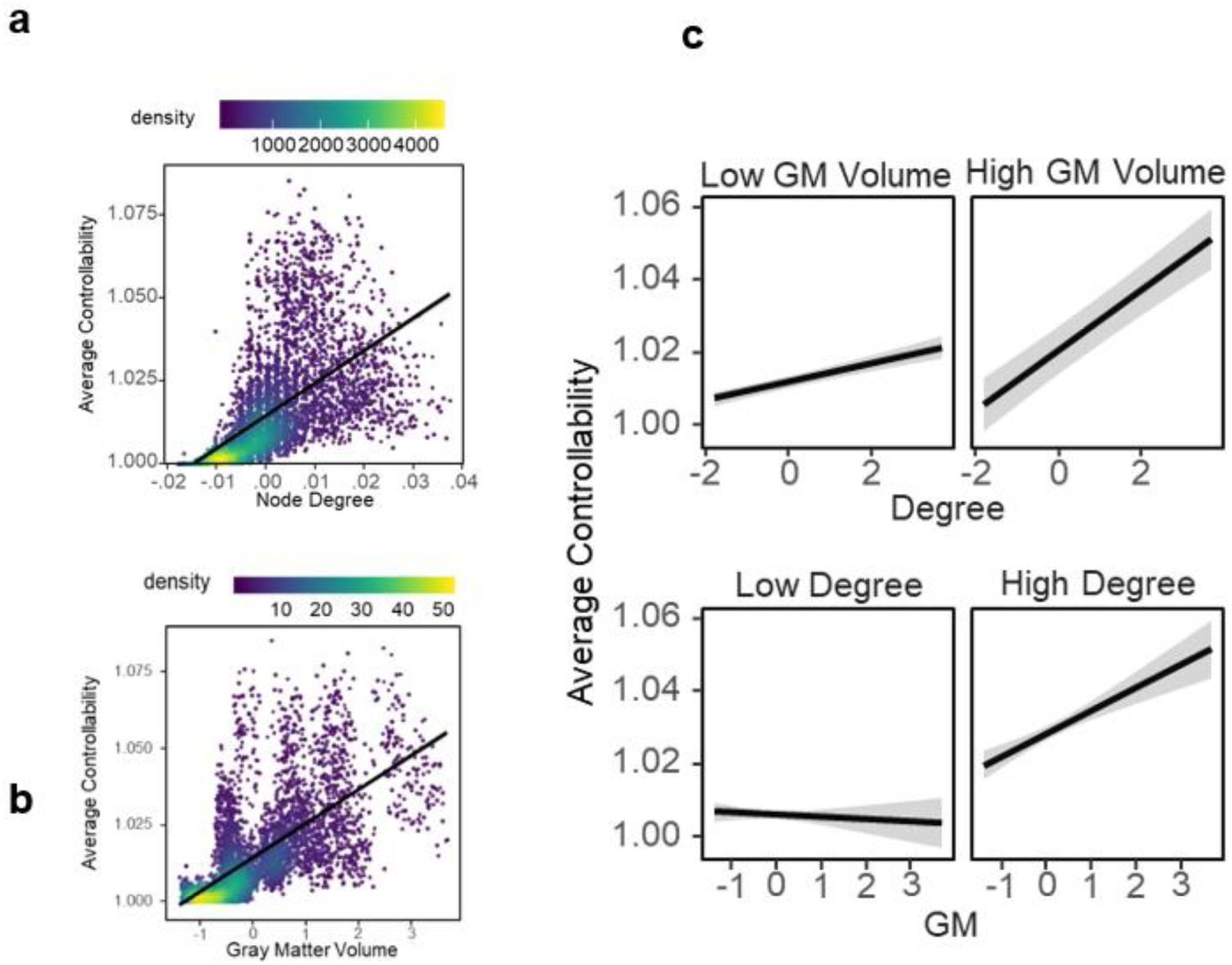
AC is estimated based on white matter structure but strongly relates to the gray matter volume. (A) Effect of degree on average controllability: mixed-effects model predicting AC corrected for regional differences of AC. In this model we corrected for GM, whole brain volume, and region. (B) Association between GM and AC. (C) Interaction between Gray Matter (GM) and Degree on AC suggesting that highest levels of AC are reached when both degree and gray matter volume are high together. For visualization, median split was used to classify GM volume and Degree into high and low respectively. In the original model, both effects were preserved as continuous variables.

In a second step we investigated if, beyond this positive association between degree and AC, GM volume explains additional variance of AC. To this aim, we extended our model by including regional GM volume and whole brain volume as additional predictors to nodal degree. Our results show, that GM volume significantly increased AC (see Figure 1B) and interacted with nodal degree (β= .42, .34 – .49, p < .001), suggesting that highest levels of average controllability were best explained with concurrent high GM volume and high node degree (see Figure 1C). Taken these results together, our results stress the interdependency between nodal connectivity strength and GM volume for brain controllability.

### 4.2 Regional distribution of Average Controllability based on gray matter volume

Further, we investigated if this global interdependency between WM and GM volume (see previous section) differs on a regional level. Our results show that, although higher GM volume and nodal degree concomitantly increased AC, there were few regions that exhibited particular sensitivity to AC based on both GM and nodal degree or only GM or nodal degree alone. Higher nodal degree in the left middle and superior frontal gyrus and left Calcarine exhibited highest AC levels, which lines up with previous research also locating driver nodes for AC in the frontal lobes. There were also regions where higher levels of nodal degree exacerbated AC, with strongest effects located in the right and left cuneus. When turning to effects of GM volume onto AC, higher volume in the right Calcarine and bilateral middle occipital gyrus exhibited higher AC levels. In contrast, higher GM volume in the right cuneus and inferior occipital gyrus exhibited lower AC levels. The finding suggests that, although on a global brain level nodal degree and GM volume concomitantly increase AC, for some regions, the effect of nodal degree counteracts with the effect of GM volume (e.g. higher nodal degree, but lower GM volume in frontal middle gyrus exhibits higher AC, and higher AC levels with higher GM volume but lower nodal degree in right cuneus).

### 4.3 Replication study

To investigate if the results in section 3.1. (higher global connectivity strength and larger GM volume increase global AC) are replicable, we used data from a replication study on a cohort of 48 subjects from another publicly available dataset where we also used a slightly different preprocessing pipeline (see Appendix for details). Also, in this data set, nodal degree and GM volume increased AC (Figure 3A and B respectively), while highest AC levels were achieved when both nodal degree and higher GM volume were high together (Figure 3C and D), suggesting that this positive association between GM volume and nodal degree is robust and not driven by individual differences in across different data sets.

**Figure 2:**
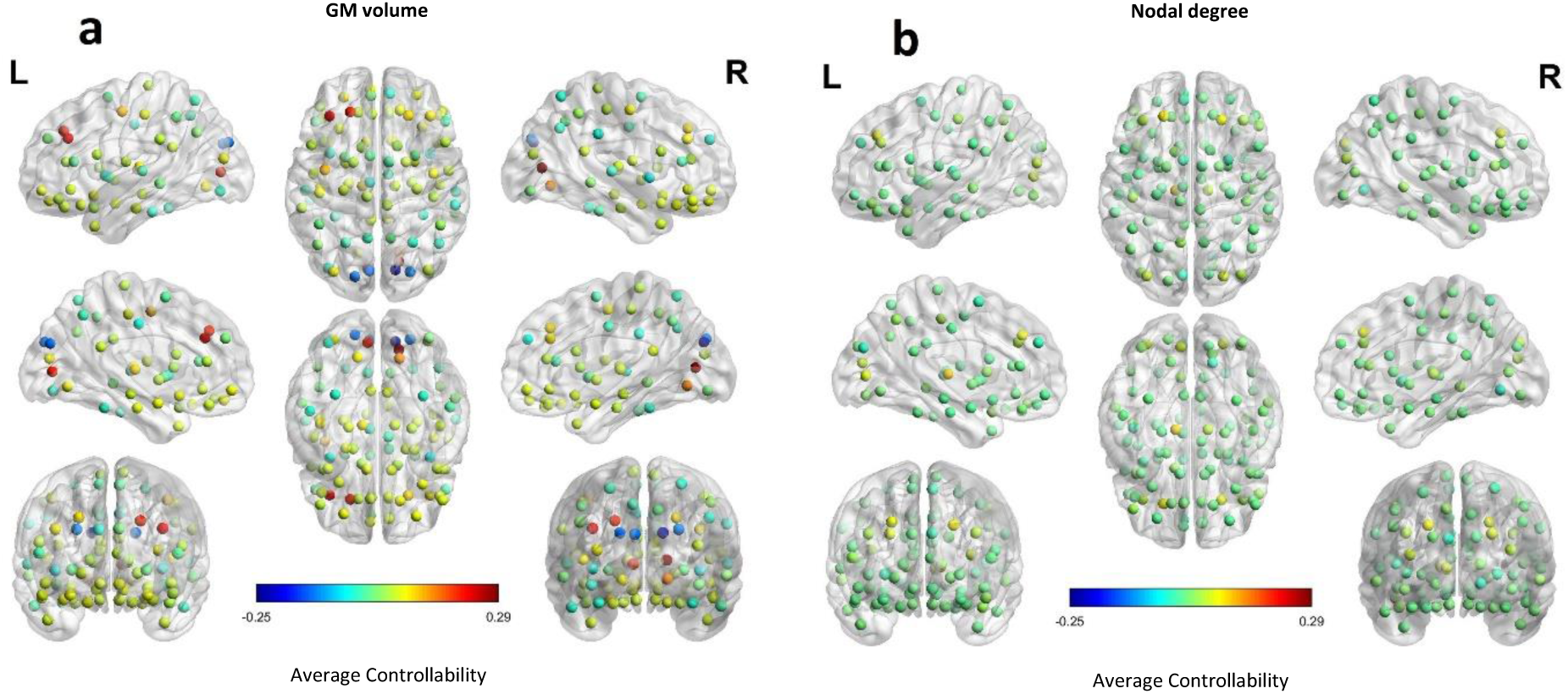
Estimated distribution of AC across regions based on gray matter volume and nodal degree onto AC derived from white matter fiber tracts. (A) and (B): regional distribution of AC based on effects of GM volume (A) and nodal degree (B). For visualization, colors represent standardized Beta coefficients for effects of GM volume and nodal degree respectively onto AC. Thus, higher values indicate a beneficial and lower values indicate an impeding effect of GM volume/nodal degree onto AC.

**Figure 3:**
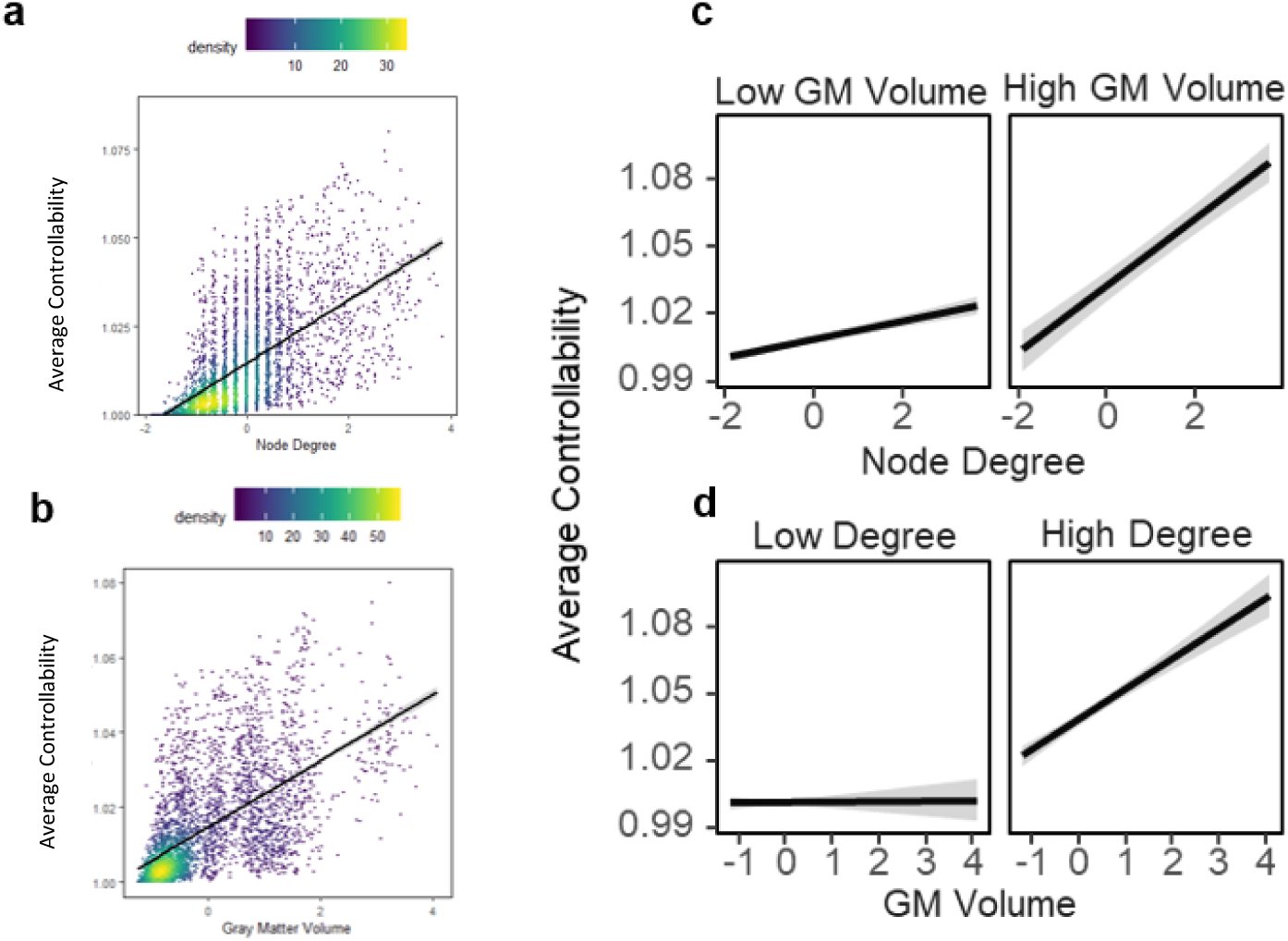
Replication sample. AC is estimated based on the connectivity of white matter fiber tracts but strongly relates to the gray matter volume. (A) Effect of nodal degree on AC. (B) Effect of GM on AC. (C) Interaction effect between Gray Matter (GM) and Degree suggesting that highest levels of AC are reached when both degree and gray matter volume are high together. For visualization, median split was used to classify GM volume and Degree into high and low respectively. In the original model, both effects were preserved as continuous variables.

## 5 Discussion

In this work, we investigated how brain volumetrics contribute to global network control properties derived from the structural connectome composed of the white matter fiber tracts. In line with (Medaglia, et al., 2017a), we hypothesized that large-scale network dynamics derived from the structural connectome (here quantified by average brain controllability) would be further explained by GM structural properties. This work is, to our knowledge, the first attempt to map the interdependency of both metrics, and we discuss findings with respect to their clinical relevance.

We show that on a global brain level, GM volume enhances the brain’s availability to dynamically transition between brain states. However, levels of brain controllability were best explained when combining metrics/information from structural properties of both WM and GM, suggesting that volumetric might provide additional information in relating brain controllability to understanding cognition, neurological and neuropsychiatric disorders, and the concept of brain reserve.

### 5.1 Mediating role of GM on the relation between WM and AC

Our finding that nodal degree is highly predictive of AC is in line with previous work (Gu, et al., 2015; Medaglia, 2019), suggesting that the brain’s ability to transverse into easy and difficult-to-reach brain states relies on sufficient strength of structural connectivity, which might reflect the degrees of freedom to steer the transition of brain states. However, our findings suggest that this picture is incomplete without the involvement of support these structural large-scale networks: on a global level, brain controllability was highest, when, in addition to high structural connectivity, GM volume was maximal. This suggests, that structural connectivity relies on sufficient support from GM reserves. A body of research highlights the relevance of whole brain volume for neuropathologic brain reserves (Bigler, 2006; Coffey, et al., 1999; Stern, 2006; Sumowski, et al., 2013), however its inter-relation with controllability metrics has been not investigated yet.

### 5.2 Potential contribution of sensory regions to brain controllability

Although our results suggest that for some of the few regions that had highest sensitivity to AC, reduced GM volume and enhanced nodal degree related to enhanced efficacy of brain controllability. This finding highlights the possible unique role of specific brain regions in brain controllability that goes against the odds of what happens globally in the brain (higher nodal degree and GM increase higher AC levels): highest effects of AC were reached with enhanced structural connectivity (nodal degree) within frontal regions, which lines up with research showing that frontal brain networks play a central role in initiating dynamic reconfigurations during executive cognition. However, increased GM volume within that very region was negatively related to brain controllability. While within clinical populations reduced GM volume is generally related to neuropathology, there is research suggesting that within healthy subjects, GM decreases with increases of WM density throughout development from adolescence to adulthood. This finding has been related to reduced quantity of synapses resulting from synaptic pruning (Giorgio, et al., 2010), which, has been predominantly found in primary visual (calcarine sulcus), and prefrontal cortex (middle frontal gyrus) (Huttenlocher, 1979; Huttenlocher and Dabholkar, 1997). In our data, brain controllability was maximal when exactly these regions showed reduced GM volume and increased connectivity of the white matter connectome. One could speculate that this finding could reflect more efficient, developmentally advanced brain functioning. On a functional level, enhanced GM volume in the right cuneus has previously been reported to predict higher error rates in a response inhibition task in bipolar (Haldane, et al., 2008) and has also been related to motor response in functional imaging studies (Booth, et al., 2005; Matthews, et al., 2005). This finding suggests that the function of those primary visual areas goes far beyond unimodal information processing.

Recent work suggests that primary sensory cortices might occupy more “hub-like” positions in the brain through enhanced long-distance connectivity across brain-wide communities (Esfahlani, et al., 2020). These findings suggest that sensory regions could be ideal hot spots for brain controllability nodes, given their high global inter-connectivity. Given that the current framework of controllability observes each brain region both as a potential controller but also controlee, it is unclear if those sensory nodes could be the controllers with respect to the afferent inputs while the other regions act as controllers for efferent demands.

### 5.3 Beyond full controllability

AC is quantified based on the eigenvalues of controllability Gramian and measures the average energy required to move the state space into all possible directions. While this is theoretically interesting and practically of relevance for potential brain stimulation techniques, more research is needed to investigate how this engineering approach translates to a brain level. Studies of dynamical functional and structural connectivity and also analysis of structural covariance have reliably shown that brain state trajectories are not random, but rather follow general rules. The theoretical significance of brain AC might overweight the practical translation of the centrality metric in analyzing the trajectories of brain states. Given that the ease of brain state transitions has different energy costs, weighted controllability metrics which are informed by the state trajectories that are experimentally observed, could significantly enhance the practical value of the analysis presented in this work.

More than the consideration discussed above, there remain more reasons why the controllability framework needs to address the partial controllable subspace. So far, the literature on applications of control theory to study brain dynamics have not sufficiently taken into account that brain regions can operate either as controllers or controllees (Medaglia, 2019). A better understanding of this association is vital to elucidate the potential role of nodes throughout large-scale dynamics based on both the white matter connectome and GM properties.

## 6 Acknowledgements

HJ was supported by Fortüne grant of Medical Faculty of University of Tübingen (No. 2487-1-0). The authors declare no conflict of interest.

## 7 Author contribution statement

Conceptualization: HJ, AZ, MW, SA. Methodology: HJ, SA, AM, AZ. Validation: HJ, AZ. Statistical analysis: AZ, HJ. Resources: MW, ME. Supervision: MW, ME, CG. Data Curation: VK, ML. Writing-Original Draft: AZ, HJ. Writing-Review: AZ, HJ, ME, MW, CG.

## 8 Data and code availability

The data used in the current study are publicly available online. See Methods for detail.

## 10 Appendix: Replication study methods

### 10.1 Data acquisition

An open-access dataset, known as ‘The Stockholm Sleepy Brain Study: Effects of Sleep Deprivation on Cognitive and Emotional Processing in Young and Old’ (see more details in (Nilsonne, et al., 2016) osf.io/bxfsb and https://openneuro.org/datasets/ds000201), was analyzed in the present study. In the original Stockholm Sleepy Brain Study, the effects of sleep deprivation on brain function with regard to emotional processing was investigated using a randomized cross-over design. The obtained anatomical, diffusion, and functional (at rest) magnetic resonance images were analyzed in the current study. After excluding the subjects with suspicious bval/bvecs tables extracted from the source DICOM files for diffusion imaging (also mentioned by the owners), 48 subjects with completed diffusion, anatomical and functional images were included in our study.

All subjects underwent diffusion MRI in a GE 3T (Discovery MR 750) scanner, using an 8-channel array head coil for signal reception and the built-in body coil for radiofrequency transmission.

Image acquisition was performed with a 3T MRI scanner (Discovery MR750, GE Medical Systems). The T1-weighted imaging with the following parameters (flip angle = 11°; TR = 64 ms; echo time TE = 2.8 ms for each image; 180 sagittal sections; spatial resolution 0.4688 × 0.46881 × 1 mm). The diffusion-weighted MRI (dMRI) acquisition protocol involved 5 non-diffusion weighted images (b-value 0 s/mm^2^) and 45 non-collinear gradient directions (b-value 1000 s/mm^2^) uniformly sampled over a sphere; *TE* = 80 ms, *TR* = 7000 ms, field of view FOV = 220 × 220 mm^2^, imaging matrix = 96 × 96, 63 consecutive slices with a thickness of 2.3 mm.

With all image files in BIDS format (https://bids.neuroimaging.io/), the quality control was performed with the help of MRIQC output prior to all processing. MRIQC is an automated processing pipeline designed to compute and compare the image quality metrics for T1 weighted anatomical and T2* weighted functional scans in both single individual level and group level (see more details in https://mriqc.readthedocs.io).

### 10.2 Anatomical data preprocessing

First, in each subject, the T1-weighted (T1w) images were corrected for intensity non-uniformity (INU) with N4BiasFieldCorrection (Tustison, et al., 2010) and then skull-stripped with the antsBrainExtraction.sh workflow (from ANTs), distributed with ANTs 2.2.0 (Avants, et al., 2008). Next, spatial normalization to the ICBM 152 Nonlinear Asymmetrical template version 2009c (Fonov, et al., 2009) was performed through nonlinear registration with antsRegistration (ANTs 2.2.0), using brain-extracted versions of both T1w volume and template. Then, brain tissue segmentation of cerebrospinal fluid (CSF), white-matter (WM) and gray-matter (GM) was performed on the brain-extracted T1w using fast (FSL 5.0.9, RRID:SCR_002823, (Zhang, et al., 2001)). Afterwards, brain surfaces were reconstructed using recon-all (FreeSurfer 6.0.1, RRID:SCR_001847, (Dale, et al., 1999). The regional volumes, defined by the parcellation scheme (Aparc+Aseg, 84 rois), were extracted from the derived output of recon-all, as well as the estimated intracranial volume. Additionally, the sub-cortical grey matter structures estimated by recon-all was replaced by the estimates from FSL’s FIRST tool.

### 10.3 Diffusion data preprocessing

The diffusion data were preprocessed by MRtrix3 (http://www.mrtrix.org/) with an adapted pipeline from the Structural connectome for Human Connectome Project (https://mrtrix.readthedocs.io/en/latest/quantitative_structural_connectivity/ismrm_hcp_tutorial.html) and Basic and Advanced Tractography with MRtrix for All Neurophiles (BATMAN, https://osf.io/fkyht/).

The mean B=0 image was calculated and used as the reference image for eddy current correction. The B1 field inhomogeneity correction for the DWI volume series and DWI data denoising by exploiting data redundancy in the PCA domain (<u>Veraart2016a</u> and <u>Veraart2016b</u>) were performed using Mrtrix3 commands: dwibiascorrect and dwidenosie, respectively. Within the estimated brain mask based on the mean b0 image, the response functions (RF) for the three different tissue types: white nd gray matter, and CSF were estimated (Dhollander and Connelly, 2016). The tissue-specific RF was further used as a kernel to perform constrained spherical deconvolution (CSD; (Tournier, et al., 2007; Tournier, et al., 2004) instead with the tensor model. Based on the derived RF for the different tissue-types, the Fiber Orientation Distribution (FOD, as explained in (Jeurissen, et al., 2014) crossing each voxel were generated (Tournier, et al., 2007; Tournier, et al., 2004). orientation distribution of the fibers in each voxel. The voxels used for RF estimation of the different tissue types was displayed on the preprocessed DW-image and visually inspected in each subject, as well as the derived FOD. Next, in order to create a whole-brain tractogram with biological plausibility of downstream streamline creation, the T1 data was registered to mean b0 image and further segmented into five tissue types for improving fiber tracking. After that, the probabilistic tractography using CSD algorithm in conjunction with Anatomically Constrained Tractography (ACT, (Smith, et al., 2012) based on the segmented five tissue types T1 was performed to create 1M streamlines in each subject. Given the bias towards streamline density of that straight track in regions of crossing fibers and the density of long tracks is overestimated by CSD algorithm, compared to short tracks, the <u>Spherical-deconvolution Informed Filtering of Tractograms (SIFT)</u> algorithm (Smith, et al., 2013) was following applied to filters the tractograms and reduce the overall streamline count to 200k. In the end, we mapped the streamlines with the above derived Aparc+Aseg as the parcellated image to produce a connectome – “structural connectivity” matrix. The edge of this matrix was defined as the count how many tracks connect each atlas region to every other of the atlas regions.

